# The weakest link: uncertainty and sensitivity analysis of extinction probability estimates for tsetse (*Glossina* spp) populations

**DOI:** 10.1101/810564

**Authors:** Elisha B. Are, John W. Hargrove

**Author notes:** Private Bag X1 Matieland Stellenbosch, 7602.

## Abstract

**Background:** A relatively simple life history allows us to derive an expression for the extinction probability of populations of tsetse, vectors of African sleeping sickness. We present the uncertainty and sensitivity analysis of extinction probability for tsetse population, to offer key insights into parameters in the control/eradication of tsetse populations.

**Methods:** We represent tsetse population growth as a branching process, and derive closed form estimates of population extinction from that model. Statistical and mathematical techniques are used to analyse the uncertainties in estimating extinction probability, and the sensitivity of the extinction probability to changes in input parameters representing the natural life history and vital dynamics of tsetse populations.

**Results:** For fixed values of input parameters, the sensitivity of extinction probability depends on the baseline parameter values. For example, extinction probability is more sensitive to the probability that a female is inseminated by a fertile male when daily pupal mortality is low, whereas the extinction probability is more sensitive to daily mortality rate for adult females when daily pupal mortality, and extinction probabilities, are high. Global uncertainty and sensitivity analysis showed that daily mortality rate for adult females has the highest impact on the extinction probability.

**Conclusions:** The strong correlation between extinction probability and daily female adult mortality gives a strong argument that control techniques to increase daily female adult mortality may be the single most effective means of ensuring eradication of tsetse population.

**Author summary:** Tsetse flies (Glossina spp) are vectors of Trypanosomiasis, a deadly disease commonly called sleeping sickness in humans and nagana in livestock. The relatively simple life history of tsetse enabled us to model its population growth as a stochastic branching process. We derived a closed-form expression for the probability that a population of tsetse goes extinct, as a function of death, birth, development and insemination rates in female tsetse. We analyzed the sensitivity of the extinction probability to the different input parameters, in a bid to identify parameters with the highest impact on extinction probability. This information can, potentially, inform policy direction for tsetse control/elimination. In all the scenarios we considered, the daily mortality rate for adult females has the greatest impact on the magnitude of extinction probability. Our findings suggest that the mortality rate in the adult females is the weakest link in tsetse life history, and this fact should be exploited in achieving tsetse population control, or even elimination.

## Introduction

Tsetse flies (*Glossina* spp) are biting flies of both public health and economic importance in 36 Sub-Saharan Africa countries. They feed exclusively on the blood of vertebrates – game animals and livestock, and also humans, and provide the link that drives the transmission of African trypanosomiasis, a tropical disease called Sleeping Sickness in humans and nagana in livestock. According to a WHO 2018 factsheet for Human Sleeping Sickness, the disease still occurs in about 36 countries in sub-Saharan Africa, mostly among poor farmers living in rural areas. Due to sustained control efforts, the number of cases of the disease has reduced. In 2015 there were about 2804 cases recorded: 97% of these were chronic infections with *Trypanosoma brucei gambiense* [1]. To sustain the reduction in cases, it is important to continue to improve understanding of the tsetse fly vector, in a bid to develop more effective control techniques: with improved cost effectiveness.

A recent study [2] employed the theory of branching processes to derive an expession for the extinction probability for closed populations of tsetse. This equation involves numerous parameters representing death, development and fertility rates during the fly’s lifecycle. These results allow us to determine, by sensitivity analysis, the relative importance of changing, through control techniques, the various parameters. Sensitivity analysis is often used to investigate the robustness of model output to parameter values [3–5]. In this context, it is important to identify the parameters that have the greatest influence on extinction probabilities of tsetse, since this information will provide insight to the eradication of tsetse, and inform policy on the direction of control efforts.

Here we adopt the model developed by Kajunguri et al [2] and Hargrove [6] for the reproductive performance of female tsetse flies inseminated by a fertile male. We then use a framework, developed by Harris [7], to derive a fixed point equation for the extinction probability for a tsetse population. This approach allows us to obtain the same expression for extinction probability as [2], but it is derived with fewer steps and with less mathematical complexity. We compute local sensitivity indices of extinction probability with respect to all input parameters by allowing the value of daily mortality rate of female pupae (*χ*) to vary between 0.001 per-day and 0.025 per-day. Due to nonlinearities and interdependencies between input parameters, local sensitivity may be highly dependent on the baseline values of the parameters [8].

To identify the most important input parameters, we use Latin Hypercube Sampling (LHS) and Partial Rank Correlation Coefficient (PRCC) methods for the global uncertainty and sensitivity of the extinction probability. The Latin Hypercube Sampling was first applied in epidemiological modelling by Blower ([9] in [10]). Several studies have since applied LHS in disease modelling, detailing its advantage over other sampling methods and describing the methodology concisely ([8]– [11]). PRCC has been used widely in determining the sensitivity of models of various systems ([8], [12], [13]) especially to assess the sensitivity of disease models to various input parameters. Combining LHS and PRCC provides a robust method for assessing the uncertainty and the sensitivity of the extinction probability to all input parameters.

In the next section, we present the branching process model developed in [2] and [6] and present an approach based on a method used in [7] to derive a fixed point equation for the extinction probability for a tsetse population. In section 3, we present the local sensitivity analysis for the extinction probability at two fixed baseline values of the input parameters and the mathematical derivation of the sensitivity indices of extinction probability w.r.t all input parameters. Section 4 presents global uncertainty and sensitivity analysis using LHS/PRCC methods. The results are discussed in detail in section 5.

## Materials and methods

The aim of this section is to develop a stochastic model for tsetse population growth in the form of a branching process and to use the model to obtain a fixed point equation for extinction probability for tsetse populations ([2], [6], [14], [15]). We develop the branching process focusing only on female tsetse flies [6]. We follow a framework developed in [6], assuming a female tsetse fly is fertilized with probability *ϵ* and survives to deposit her first larva with probability *λ*^*ν*+*τ*^ : *ν* is days to first ovulation, *τ* is the inter-larval period, and *λ* is adult female daily survival probability. She produces a female pupa with probability *β*, and the pupa survives to adulthood with probability *ϕ*^*g*^ (where *g* is the pupal duration and *ϕ* is the daily survival probability of the pupa). The mother dies before the next pregnancy, having produced a single surviving daughter, with probability (1− *λ*^*τ*^). The probability that an adult female tsetse dies after producing a single surviving daughter after surviving one pregnancy is:

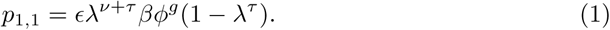

Equation (1) can be generalized by induction to obtain the probability that a female tsetse produces *k* surviving female offspring after surviving *n* pregnancies. Thus

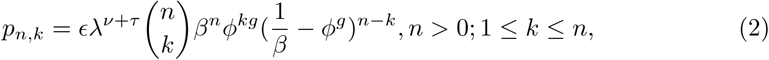

where 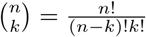 is the binomial coefficient.

Suppose *p*_0_, *p*_1_, *p*_2_, … are the probabilities that a female tsetse produces 0, 1, 2, … surviving female offspring in her lifetime, respectively. Suppose also that *p*_0_ + *p*_1_ *<* 1, to avoid the trivial case where a tsetse fly only produces 0 or 1 female offspring.

Summing Equation (2) over all *n*, gives *p*_*k*_, the probability that a female produces *k* surviving female offspring in its lifetime.

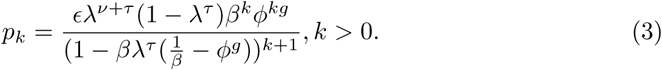

Equation (3) was used in [2] to obtain the mean and variance of the population size, extinction probability and time to extinction of populations of tsetse. Proofs of equations (1) and (2) are provided in [2] (Supplementary Information).

It can be shown easily that *p*_0_, *p*_1_, *p*_2_, … follow a geometric series, such that *p*_*k*_ = *bc*^*k*−1^, *k* = 1, 2, 3, …, where *b, c >* 0; and 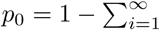. Equation (3) then becomes:

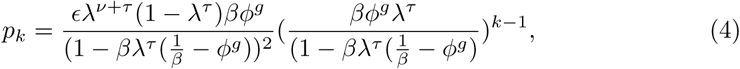

where 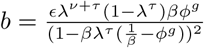 and 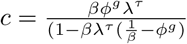.

Following a framework developed by Harris [7], the generating function *g*(θ) of *p*_*k*_, is a fractional linear function given by;

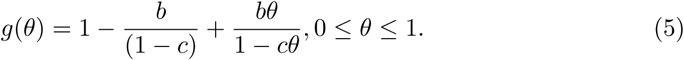

### Extinction probability

The extinction probability for tsetse population is the non-negative fixed point of Equation (5), i.e. 0 *≤θ ≤*1 such that *g*(*θ*) = *θ*.

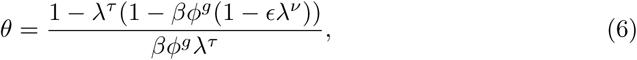

where *β ϕ*^*g*^*λ*^*τ*^ ≠ 0. In practice, 0 *< β <* 1, 0 *< λ <* 1 and 0 *< ϕ <* 1. This implies that the survival probabilities for both adult females and female pupa, and the probability that a pupa deposited is female are all in the open interval (0, 1). Which allows us to avoid a case where *θ*=0 or 1 trivially. Equation (6) is the solution for the situation where the initial population consists of just a single female fly. For *N* flies in the pioneer population, and assuming that the survival and reproductive rates of all individual flies are independent, extinction probability is *θ*^*N*^.

### Local sensitivity analysis of *θ*

In this section, we perform local sensitivity analysis, otherwise known as elasticity analysis, on the extinction probability for tsetse populations. Given that the extinction probability *θ*, depends differentiably on each input parameter, the normalized forward sensitivity (elasticity) index of *θ* w.r.t all input parameters is:

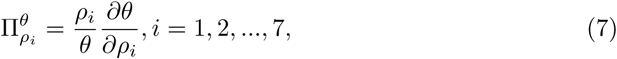

where *ρ*_*i*_ is the set of all input parameters of the extinction probability. This method has been used extensively in the literature to determine the sensitivity of the reproduction number *R*_*o*_ of epidemiological models to model parameters [4, 5, 16]. When the initial population consists of *N* female tsetse, the extinction probability is *θ*^*N*^. The sensitivity indices of *θ*^*N*^ w.r.t all input parameters is;

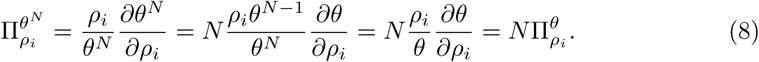

Notice that, when there are *N* female flies in the initial population, the sensitivity indices of *θ*^*N*^ w.r.t all input parameters is the sensitivity indices of *θ* multiplied by *N*. The larger the size of the initial population, the more sensitive extinction probability is to input parameters.

Writing equation (6) in terms of daily mortality rate for adult females (*ψ*), and the daily mortality rate for female pupae (*χ*), yields:

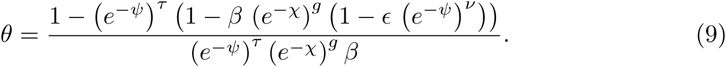

Table 1 shows the derivations of the sensitivity indices of extinction probability with respect to all seven input parameters. These expressions were derived from equations (7) and (9) with a simple code in MAPLE 17 environment.

**Table 1.** Expressions for the sensitivity indices of extinction probability with respect to all seven input parameters.

| Parameters | The sensitivity of extinction probability ($\theta$) to input parameters |
| --- | --- |
| $\psi$ | $\Pi_{\beta}^{\theta} = -\frac{-1+(e^{-\psi})^{\tau}}{-1+(e^{-\psi})^{\tau}-(e^{-\psi})^{\tau}(e^{-\chi})^g\beta+(e^{-\psi})^{\tau+\nu}(e^{-\chi})^g\beta\epsilon}$ |
| $\chi$ | $\Pi_{\chi}^{\theta} = \frac{\chi g(-1+(e^{-\psi})^{\tau})}{-1+(e^{-\psi})^{\tau}-(e^{-\psi})^{\tau}(e^{-\chi})^g\beta+(e^{-\psi})^{\tau+\nu}(e^{-\chi})^g\beta\epsilon}$ |
| $\epsilon$ | $\Pi_{\epsilon}^{\theta} = \frac{(e^{-\psi})^{\tau+\nu}(e^{-\chi})^g\beta\epsilon}{-1+(e^{-\psi})^{\tau}-(e^{-\psi})^{\tau}(e^{-\chi})^g\beta+(e^{-\psi})^{\tau+\nu}(e^{-\chi})^g\beta\epsilon}.$ |
| $g$ | $\Pi_{\chi}^{\theta} = \frac{\chi g(-1+(e^{-\psi})^{\tau})}{-1+(e^{-\psi})^{\tau}-(e^{-\psi})^{\tau}(e^{-\chi})^g\beta+(e^{-\psi})^{\tau+\nu}(e^{-\chi})^g\beta\epsilon}$ |
| $\psi$ | $\Pi_{\psi}^{\theta} = -\frac{\psi((e^{-\psi})^{\tau+\nu}(e^{-\chi})^g\beta\epsilon\nu+\tau)}{-1+(e^{-\psi})^{\tau}-(e^{-\psi})^{\tau}(e^{-\chi})^g\beta+(e^{-\psi})^{\tau+\nu}(e^{-\chi})^g\beta\epsilon}.$ |
| $\tau$ | $\Pi_{\tau}^{\theta} = \frac{\tau \ln(e^{-\psi})}{-1+(e^{-\psi})^{\tau}-(e^{-\psi})^{\tau}(e^{-\chi})^g\beta+(e^{-\psi})^{\tau+\nu}(e^{-\chi})^g\beta\epsilon}.$ |
| $\nu$ | $\Pi_{\nu}^{\theta} = \frac{\nu \beta (e^{-\chi})^g (e^{-\psi})^{\tau+\nu} \epsilon \ln(e^{-\psi})}{-1+(e^{-\psi})^{\tau}-(e^{-\psi})^{\tau}(e^{-\chi})^g\beta+(e^{-\psi})^{\tau+\nu}(e^{-\chi})^g\beta\epsilon}.$ |

## Results

Table 2 shows the sensitivity indices of extinction probability w.r.t all input parameters for different values of extinction probabilities. For instance, the sensitivity indices of *θ* to *ϵ* (probability female is inseminated by a fertile male) decreases by *>* 60% when *θ* (extinction probability) approaches 1, implying that, at *θ* = 0.419, a 10% decrease in *ϵ* will yield a 22% increase in *θ*, whereas, at *θ* = 0.96, a 10% decrease in *ϵ* will only yield an 8.7% increase in *θ*.

**Table 2.** List and description of parameters affecting extinction probabilities for tsetse populations, and the sensitivity indices for these parameters, at two different values of extinction probability.

| Parameters & descriptions | Baseline values | Sensitivity indices |  |
| --- | --- | --- | --- |
| | | $\theta = 0.419$ | $\theta = 0.960$ |
| Daily mortality rate for adult females ($\psi = -\ln(\lambda)$) | 0.02-0.03 per-day [17] | +1.030 | +1.080 |
| Daily mortality rate for female pupae ($\chi = -\ln(\phi)$) | 0.01-0.025 per-day [17] | +0.507 | +0.374 |
| Probability deposited pupa is female ($\beta$) | 0.5 [6] | -0.836 | -0.832 |
| Probability female is inseminated by a fertile male ($\epsilon$) | 1 [6] | -2.220 | -0.870 |
| Inter-larval period ($\tau$) | 9 days [6] | +0.875 | +0.929 |
| Pupal duration ($g$) | 27 days [6] | +0.507 | +0.374 |
| Time from adult female emergence to first ovulation ($\nu$) | 7 days [6] | +0.158 | +0.154 |

### Varying sensitivity indices of *θ* w.r.t all input parameters as a function of *χ*

Here we investigate the changes that occur in the sensitivity indices of extinction probability with respect to six input parameters by allowing *χ* to vary between 0.1% to 2.5%. A simple script was written in MAPLE 17 environment to calculate the local sensitivity indices of *θ* w.r.t to the six remaining input parameters for different values of *χ*. Figure 1(A and B) show changes in the sensitivity indices of *θ* w.r.t to each parameter as the daily mortality rate for female pupae (*χ*) varies from 0.1% to 2.5%, while keeping constant the other baseline values of *g, τ, ν, Ψ, β*, and *ϵ* (Table 1).

**Fig 1.**
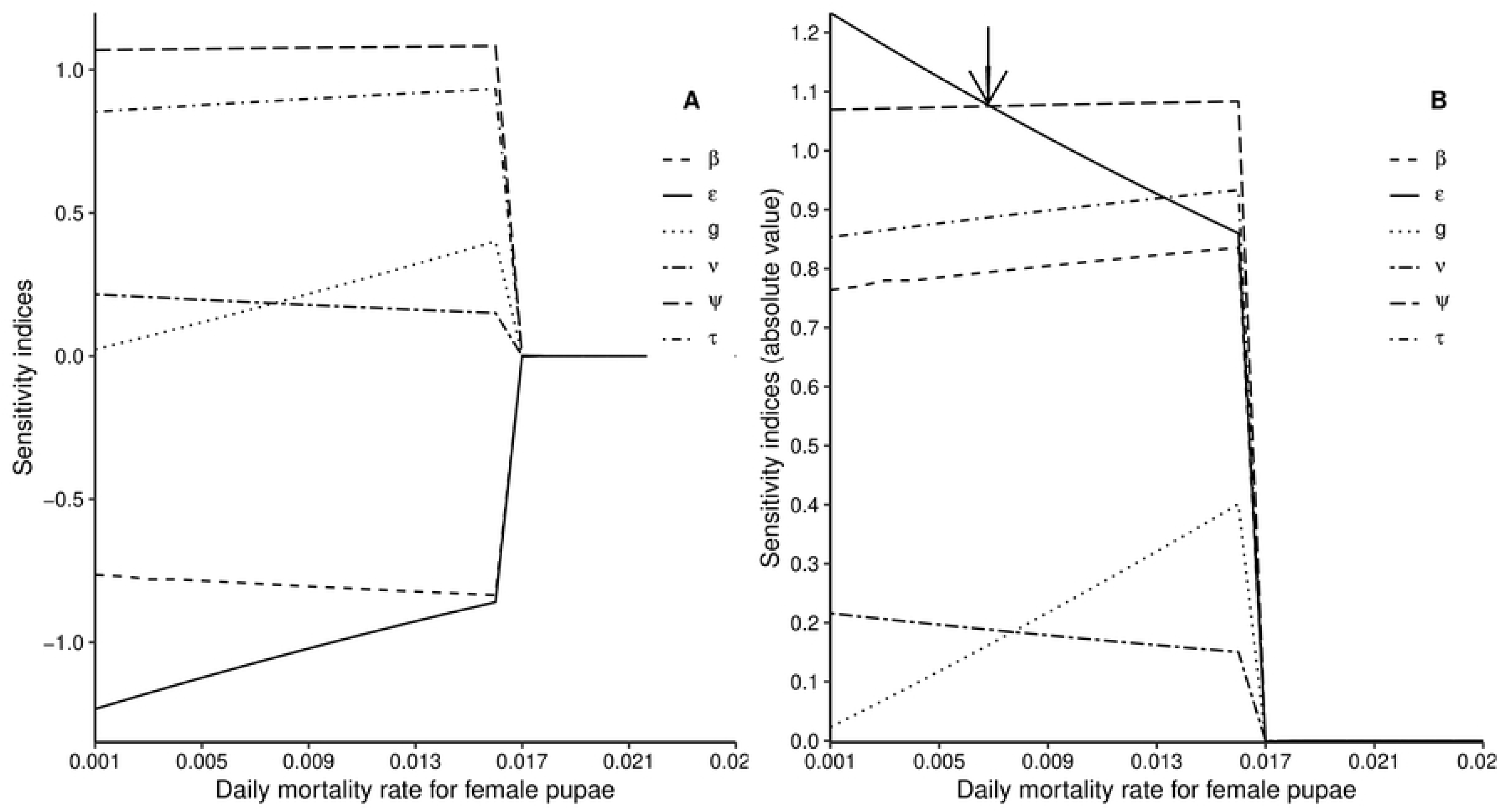
Variation in the sensitivity of extinction probability *θ* to six input parameters (*β, ϵ, ν, g, Ψ, τ*) as a function of the values of the background daily rate (*χ*) of pupal female mortality. (A) The sensitivity indices of extinction probability to six input parameters with signs. (B) The sensitivity indices of extinction probability to six input parameters, in absolute value. The arrow through B indicates the point where *θ* becomes more sensitive to *Ψ* than *ϵ*.

As *χ* increases from 0.001 to 0.0065, the sensitivity index of *θ* w.r.t *ϵ* reduces below the sensitivity index of *θ* w.r.t *Ψ*. At that point extinction probability becomes more sensitive to *Ψ* than *ϵ*. When *χ* increases further to 0.013, the sensitivity of extinction probability to *ϵ* drops further below the sensitivity of extinction probability to *τ* (Fig 1 (A and B)).

Local sensitivity analysis may not be robust enough to capture the actual influence of all input parameter values on the extinction probability since there are interdependencies between input parameters. We, therefore, proceed to carry out global uncertainty and sensitivity analysis of the extinction probability for tsetse population.

### Global uncertainty and sensitivity analysis of *θ*

The exact values of the input parameters are not known in field suituatins, where many of these parameters depend on temperature and other climatic factors. It is therefore important to quantify the uncertainty involved in estimating the extinction probability. To quantify the uncertainty involved in estimating the extinction probability (*θ*) and to establish the most important input parameters, we use LHS and PRCC methods for the global uncertainty and sensitivity analysis of the extinction probability. The method follows the approach of Samsuzzoha et al [4].

### Uncertainty analysis

We aim to analyse the uncertainty involved in quantifying extinction probability (*θ*) based on the uncertainties associated with the input parameters. Accordingly, in order to investigate the sensitivity to this uncertainty we sample values from prior distributions of these parametes. We define prior probability distribution functions for each of the input parameters, based on the studies done on the life cycle of tsetse published in the literature [17, 18]. The probability distribution functions are given in Table 3, where *β, N* and *U* denote beta, normal and uniform distributions, respectively.

**Table 3.**
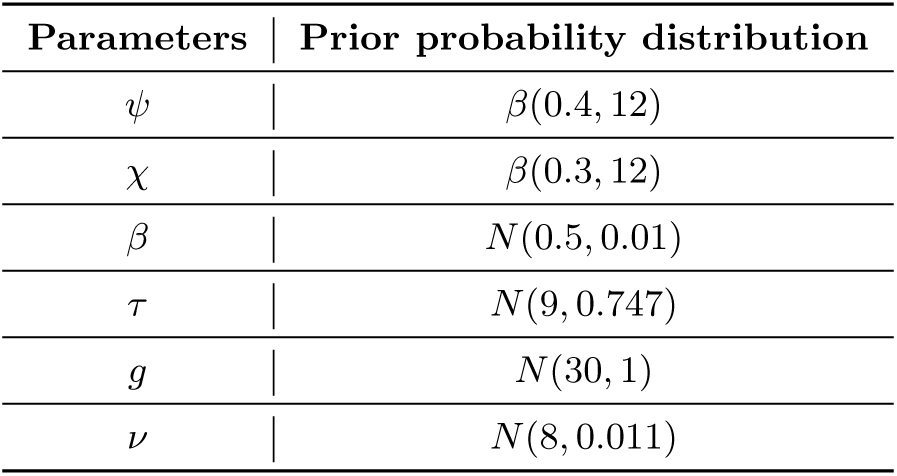
List of parameters and their prior probability distributions.

| Parameters | Prior probability distribution |
| --- | --- |
| $\psi$ | $\beta(0.4, 12)$ |
| $\chi$ | $\beta(0.3, 12)$ |
| $\beta$ | $N(0.5, 0.01)$ |
| $\tau$ | $N(9, 0.747)$ |
| $g$ | $N(30, 1)$ |
| $\nu$ | $N(8, 0.011)$ |

Using LHS, we obtain the uncertainty output for all the input parameters and also for the extinction probability. LHS is used to sample from the stratified probability distribution functions for different parameters. Using 1000 intervals of equal probabilities. Figure 2 shows the uncertainty output for all the input parameters and the shape of their probability distribution together with their summary statistics. The uncertainty output for extinction probability (*θ*) shows that it is beta distributed with mean = 0.415 and standard deviation = 0.386.

**Fig 2.**
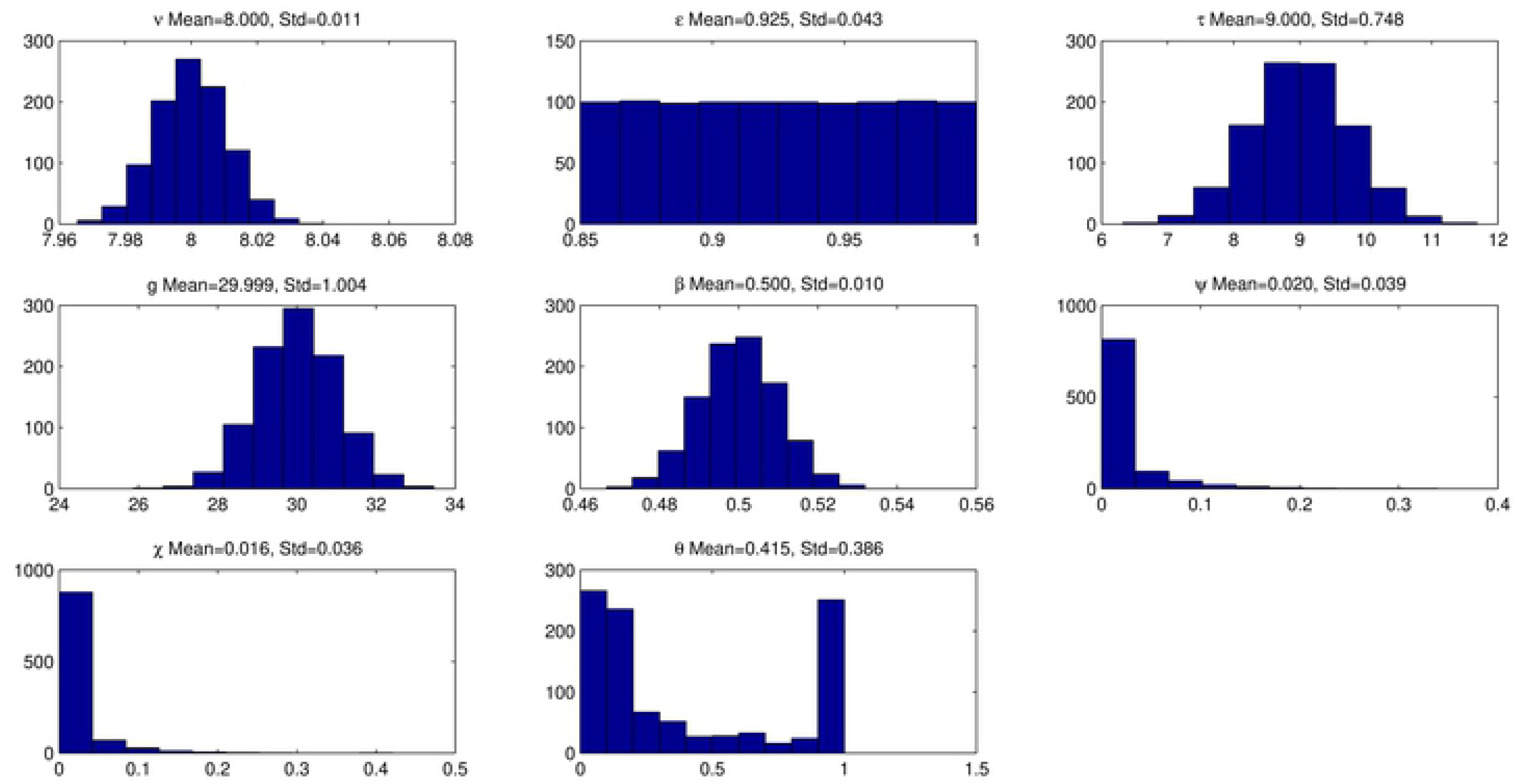
The uncertainty output for all input parameters, together with uncertainty output of the extinction probability, obtained from Latin hypercube sampling using a sample size of 1000 for the seven input parameters. Each parameter appears at the top of the corresponding sub-plot.

### PRCC/sensitivity indices of *θ* w.r.t all input parameters

To identify key input parameters, we carry out a sensitivity analysis by calculating the PRCC between each input parameter and the extinction probability. The parameter with the highest PRCC has the largest influence on the magnitude of the extinction probability. Figure 3 shows the PRCC outputs for all input parameters, where the probability (*ϵ*) that a female fly is inseminated by a fertile male is essentially equal to 1. In the field, males manage to find and mate with females, even when population levels are quite low [19]. For most tsetse populations, the probability of insemination is thus close to 1. Accordingly, we allow *ϵ* to vary between 0.999-1. In figure 5, the prior probability distributions are kept the same save for *ϵ* which is sampled between 0.885 and 1 [10, 11, 20].

**Fig 3.**
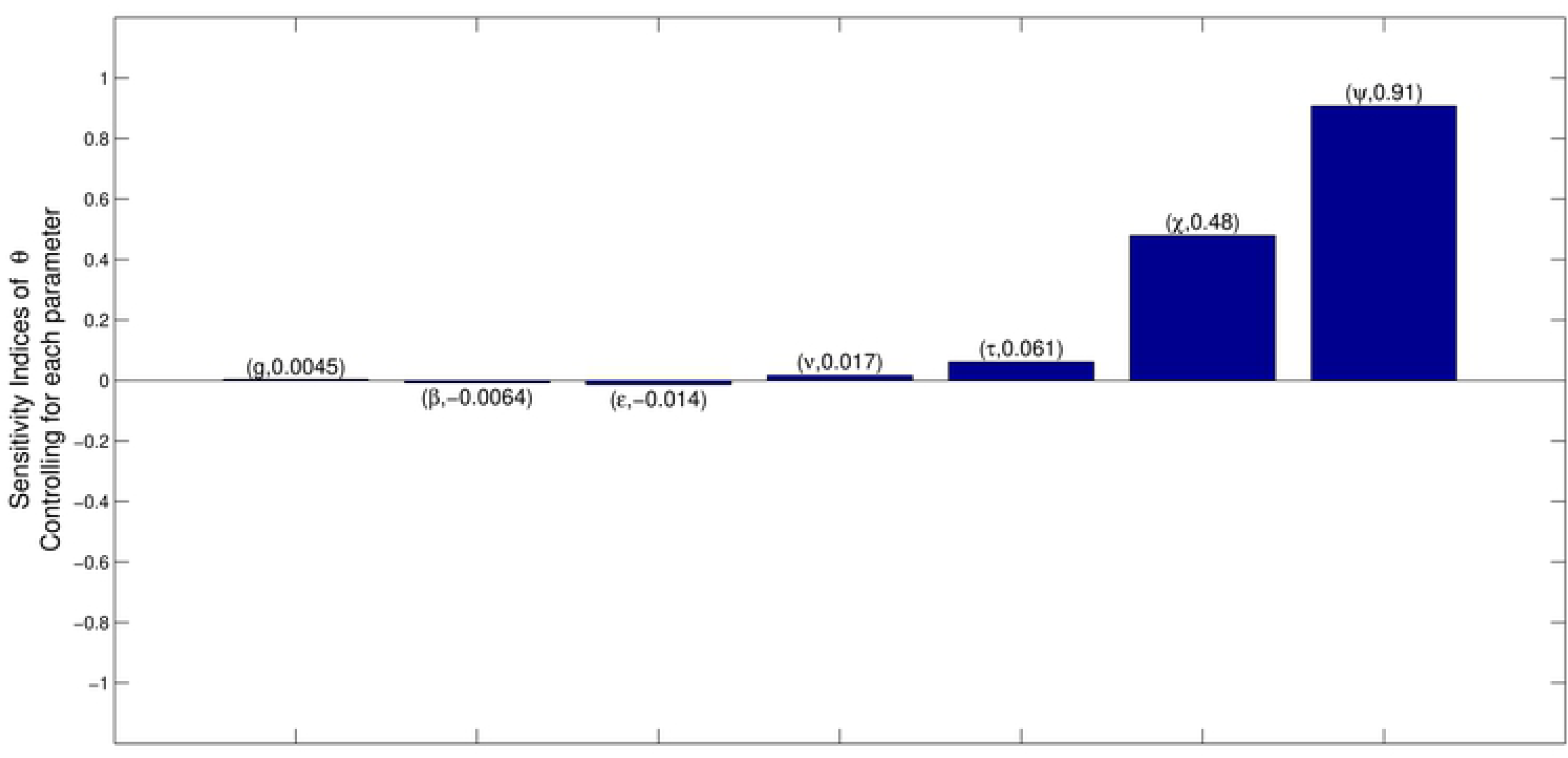
PRCC output for all input parameters with respect to the extinction probability. Sampling *ϵ* between 0.999 and 1.

**Fig 4.**
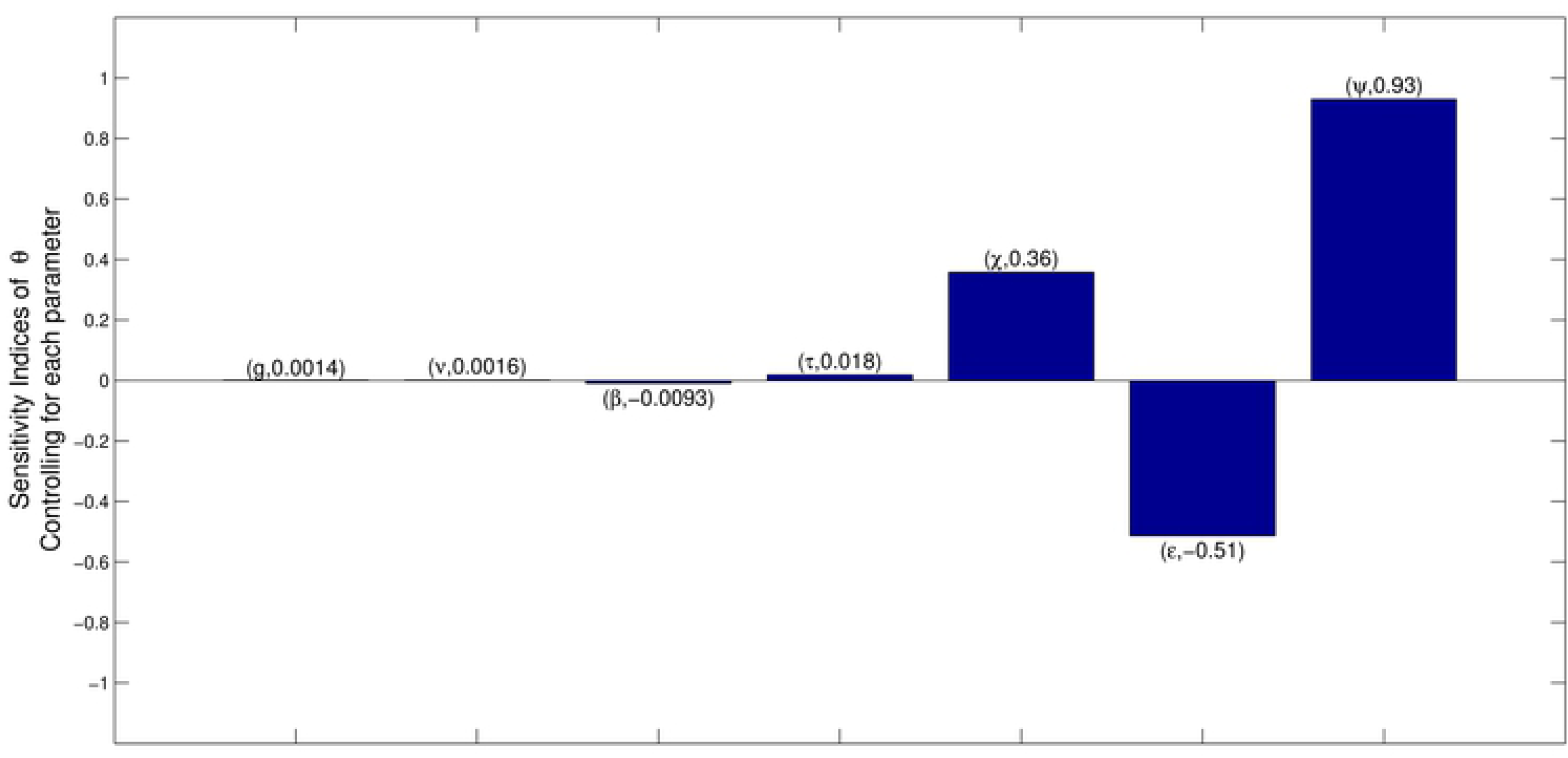
PRCC output for all input parameters with respect to the extinction probability. Sampling *ϵ* between 0.855 and 1

**Fig 5.**
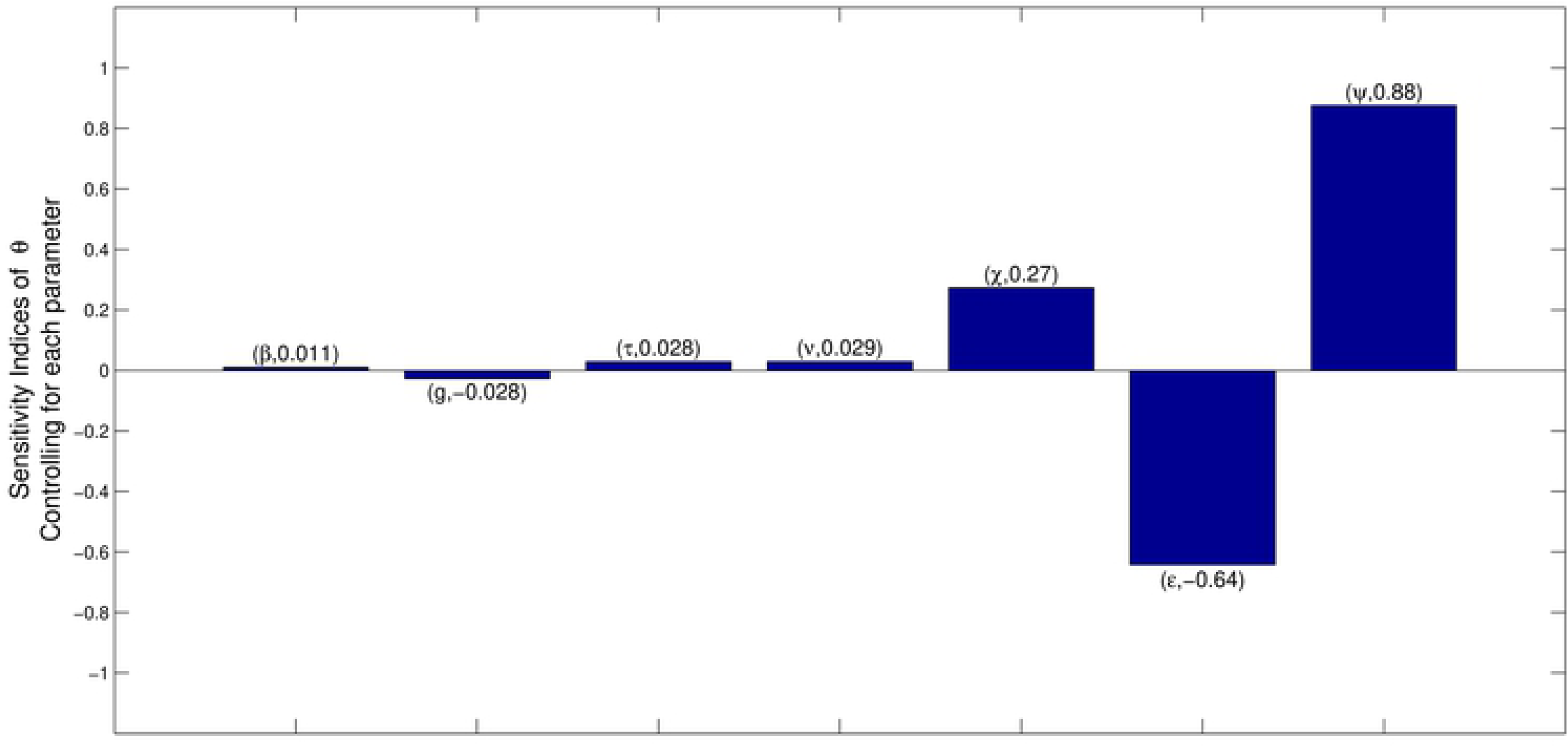
PRCC output for all input parameters with respect to the extinction probability. Sampling *ϵ* between 0.51 and 1

LHS is used to sample from the prior probability distributions, where *ϵ* is sampled from a uniform distribution *U* (0.999, 1). Figure 3 shows that daily mortality rate for adult females (*Ψ*) has a strong correlation with the extinction probability with PRCC score 0.91, followed by daily mortality rate for female pupae (*χ*) and inter-larval period (days) (*τ*) having PRCC scores 0.47 and 0.058, respectively.

The female tsetse fly generally mates only once in her lifetime, storing the sperm in spermathecae and using small amounts to fertilize her eggs one at a time [21, 22]. When sterile males are introduced into tsetse population, the probability (*ϵ*) that a female is inseminated by a fertile male falls below unity, by an amount that depends on the ratio of sterile to fertile males in the population.

The Sterile Insect Technique (SIT) has been used in attempts to control tsetse flies populations [23, 24] and was used to eradicate a small population of *G*. *austeni* on Unguja Island, Zanzibar, Tanzania [25]. The probability that a female is inseminated by a sterile male is 1 − *ϵ*. We now allow baseline values of *ϵ* to vary over a wide range, in order to assess the sensitivity of extinction probability to changes in *ϵ* at varying baseline levels of the proportions of sterile males in the population. Figures 4 – 6 show the PRCC scores when *ϵ* is uniformly distributed either as U(0.855, 1), U(0.51, 1) or U(0.1, 1). The PRCC scores for *ϵ* in these three scenarios were −0.51, −0.64 and −0.72, respectively. Thus the absolute value of the PRCC score for *ϵ* increases as we allow more variability in the probability distribution function.

**Fig 6.**
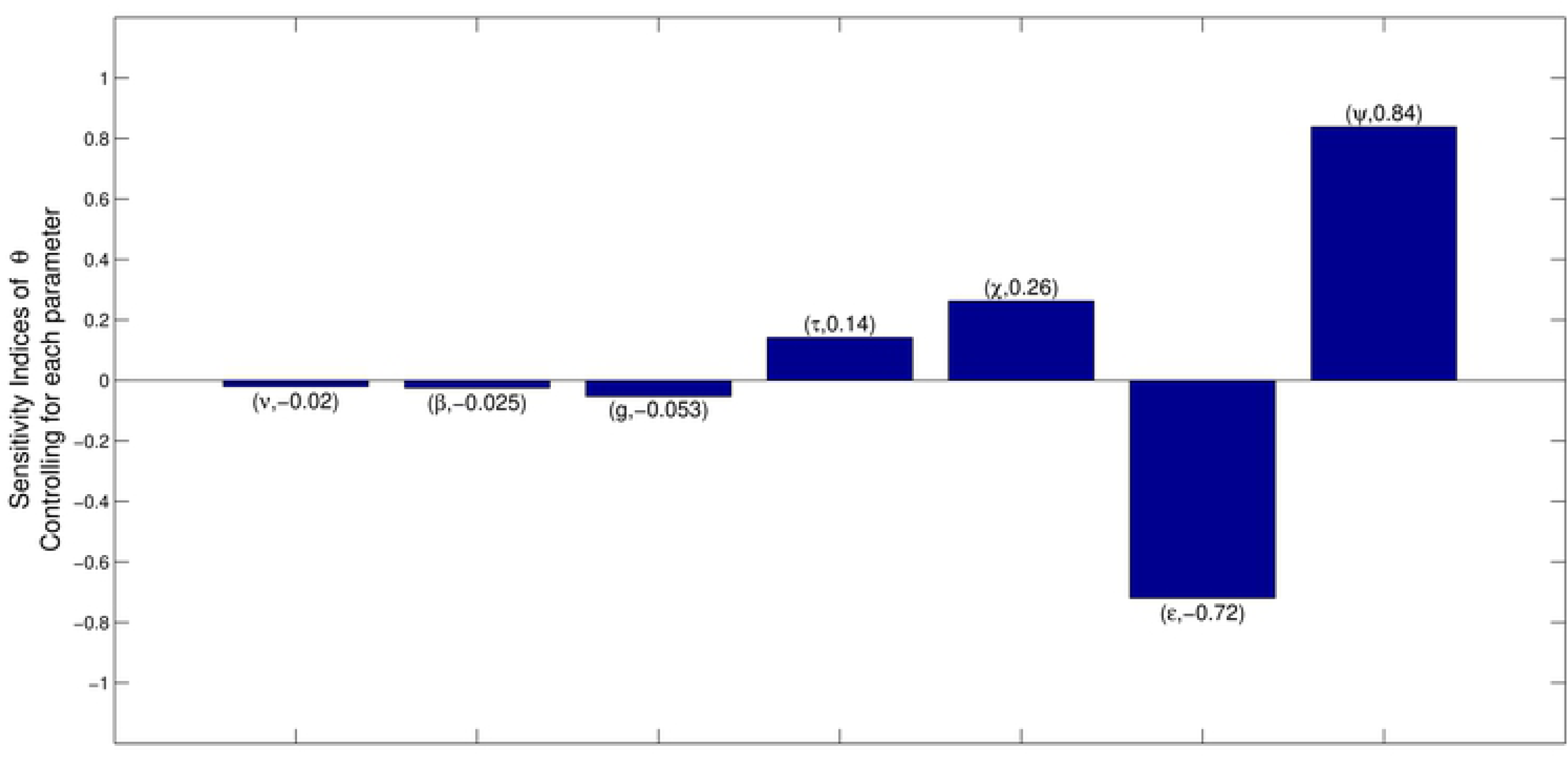
PRCC output for all input parameters with respect to the extinction probability. Sampling *ϵ* between 0.1 and 1

## Discussion

The simple life history of the tsetse fly enabled us to model its population dynamics as a stochastic branching process. We derived an expression for the extinction probability for tsetse populations and performed local and global sensitivity analyses, as well as global uncertainty analysis, on the extinction probability. We calculated all results for two fixed baseline values for *χ*, corresponding to values that resulted in low or high extinction probabilities. We obtained the sensitivity indices of the extinction probability to seven input parameters. When the extinction probability (*θ*) is fixed at either low or high levels (0.419 or 0.960) *θ* is more sensitive to changes in daily adult mortality (*Ψ*) and the fertile insemination probability (*ϵ*) than to any of the other parameters. For a change in *θ* from 0.419 to 0.960, the sensitivity index of *θ* w.r.t. *Ψ* increases by 0.05, whereas the change w.r.t. *ϵ* is larger, at 1.35 (decrease in absolute value) (Table2). The parameters *Ψ* and *ϵ* are important as they underpin the two main approaches to tsetse control. Hocking et al [26] broadly classified tsetse control and elimination techniques to include: game destruction, bush clearing, use of insecticides and biological control. These techniques can be pooled into two fundamental control philosophies - those which aim, primarily, to increase mortality rates in adult flies and those, like SIT, which aim to reduce tsetse birth rates [24]. Sensitivity analysis will indicate which parameter out of the two has the highest impact on the extinction probability.

From Table 2, observe that the sensitivity indices of *θ* to the input parameters depends on the value of the extinction probability. We allowed the daily mortality rate for pupae (*χ*) to vary from 0.001 to 0.025. The lower and upper bound values result in low and high extinction probabilities, respectively. We then calculated the sensitivity indices of *θ* w.r.t. the remaining six parameters. Figure 1(A and B) shows that the sensitivity of *θ* to each of the input parameters changes as extinction probability increases with increasing values of *χ*. Observe that for *χ ≥* 0.018, the sensitivity indices of all the six parameters converged to zero. This is expected since the set baseline parameters values for all input parameter will correspond to extinction probability (*θ*) = 1 at *χ ≥* 0.018. This can be verified easily, by substituting parameter values into equation (6).

LHS and PRCC provide a suitable technique for assessing the impact of input parameters on the output and therefore inform possible choices for effective control efforts [27]. We defined prior probability density functions for the seven input parameters and we sampled from intervals of equal probability using LHS. The PRCC score of all input parameters was obtained for three sets of the probability distribution function, fixed for six parameters and varied only for *ϵ*. In all cases, *Ψ* has the strongest impact on the extinction probability. The PRCC score for *ϵ* increases as we allow for more variability in its prior probability distribution.

The SIT is an effective technique used to suppress or eradicate populations of tsetse, but its major drawback is the large number of sterile flies that have to be produced and introduced into the wild [23]. Our results confirm this; the higher the number of sterile males introduced into the wild, the higher the impact of *ϵ* on the extinction probability (Figs 3–6).

## Conclusions

In all scenarios considered, control techniques which can achieve high mortality rates for adult female flies have the strongest impact on extinction probability. Control techniques such as SIT, which can reduce reproductive rates, without increasing mortality, can also have a strong impact on extinction probability. This happens only when the number of sterile males, introduced into the population, massively outnumber wild males, such that the probability is low that a virgin female will mate with a fertile male.

A limitation of our work is the assumption that the tsetse flies experience fixed environmental conditions throughout their life history. This assumption is not true in the wild, where tsetse experience daily and seasonal changes in various climatic effects. We will address this problem in future work.

